# Activity of Selected Nucleoside Analogue ProTides against Zika Virus in Human Neural Stem Cells

**DOI:** 10.1101/533497

**Authors:** Jean A. Bernatchez, Michael Coste, Sungjun Beck, Grace A. Wells, Lucas A. Luna, Alex E. Clark, Zhe Zhu, Christal D. Sohl, Byron W. Purse, Jair L. Siqueira-Neto

## Abstract

Zika virus (ZIKV), an emerging flavivirus which causes neurodevelopmental impairment to fetuses and has been linked to Guillain-Barré syndrome, continues to threaten global health due to the absence of targeted prophylaxis or treatment. Nucleoside analogues are good examples of efficient anti-viral inhibitors, and prodrug strategies using phosphate masking groups (ProTides) have been employed to improve the bioavailability of ribonucleoside analogues. Here, we synthesized and tested a library of 13 ProTides against ZIKV in human neural stem cells. Strong activity was observed for 2′-*C*-methyluridine and 2′-*C*-ethynyluridine ProTides with an aryloxyl phosphoramidate masking group. Conversion of the aryloxyl phosphoramidate ProTide group of 2′-*C*-methyluridine to a 2-(methylthio)ethyl phosphoramidate completely abolished antiviral activity of the compound. The aryloxyl phosphoramidate ProTide of 2′-*C*-methyluridine outperformed the hepatitis C virus (HCV) drug sofosbuvir in suppression of viral titers and protection from cytopathic effect, while the former compound’s triphosphate active metabolite was better incorporated by purified ZIKV NS5 polymerase over time. Molecular superpositioning revealed different orientations of residues opposite the 2′-fluoro group of sofosbuvir. These findings suggest both a nucleobase and ProTide group bias for the anti-ZIKV activity of nucleoside analogue ProTides in a disease-relevant cell model.

## 1. Introduction

The explosive spread of Zika virus (ZIKV) during the 2015-2016 epidemics in Latin America attracted worldwide attention to this previously neglected disease. The lack of effective vaccines or small molecules to prevent or treat this infection remains a cause for concern and emphasizes the urgent need for new therapeutic options [1]. ZIKV, an emerging flavivirus infection, causes several serious neurodevelopmental anomalies in the fetus, including microcephaly, congenital ZIKV syndrome (CZS), and even fetal demise [2]. While most cases of ZIKV infection are asymptomatic, reports of rash, conjunctivitis, pain in the muscles or joints, and fever have been recorded in clinical manifestations of the disease, and in rare cases, ZIKV infection has been linked to the neuroinflammatory Guillain-Barré syndrome [2].

Viral polymerases remain attractive drug targets for the development of selective antiviral therapies [3–5]. Generally, clinically-approved inhibitors that target these proteins fall into two broad classes [6,7]. The first class consists of nucleoside analogues that mimic the natural substrate of the enzyme. Upon analogue incorporation by the virally-encoded polymerase, DNA or RNA synthesis is abrogated by preventing further nucleotide incorporation (chain termination), thereby arresting viral replication. The second class is known as non-nucleoside inhibitors, which bind allosterically and arrest viral nucleic acid synthesis by distorting the polymerase active site geometry to interfere with nucleotide binding or nucleotide incorporation.

A common prodrug strategy used for antiviral ribonucleoside analogues involves the chemical synthesis of nucleoside analogue monophosphates with metabolically-removable masking groups [8–10]. These masking groups neutralize the negative charge of the phosphate on the nucleoside and allow for better membrane penetrance of these compounds. Chemical addition of the first phosphate group is crucial for improving intracellular levels of the active triphosphate metabolite form of ribonucleoside analogues. Metabolic addition of the first phosphate group is relatively slow for this class of compounds, and represents a rate-limiting step for the triphosphorylation of ribonucleoside analogues to active drug molecules.

This *“*ProTide*“* strategy has been successfully deployed in the synthesis of bioactive ribonucleoside analogues [11,12], including the FDA-approved hepatitis C virus (HCV) drug sofosbuvir [13]. Sofosbuvir has recently been shown by multiple groups to also be active against ZIKV *in vitro* and *in vivo* [14–17], demonstrating that ribonucleoside analogue ProTides are an attractive avenue for the development of novel, selective antivirals against ZIKV.

Numerous nucleoside analogues have recently been explored as antiviral agents against ZIKV [18–23]. However, many of these compounds were tested in either cell-free systems or using cells that are not clinically relevant for the disease (such as Vero cells). Further, sofosbuvir is the only inhibitor tested against ZIKV thus far in ProTide form. In addition, recent work by our group and others has suggested that sofosbuvir and ProTides in general have differential activity depending on the cell line used, which may be linked to the cell-specific metabolism of ProTides[17,24]. In this work, we chemically synthesized a library of 13 ProTides and tested them for activity against ZIKV in human neural stem cells, a disease-specific cell model of infection. Interestingly, the activity of 2′-*C*-modified aryloxyl phosphoramidate ProTides was strongly biased towards uridylate derivatives, with only modest activity observed for the 2′-*C*-modified adenylate and cytidylate aryloxyl phosphoramidate ProTides we tested. Changing the aryloxyl phosphoramidate masking group to a 2-(methylthio)ethyl phosphoramidate abolished the activity of the 2′-*C*-methyluridine ProTide, a representative hit from our library. In a head-to-head comparison between the 2′-*C*-methyluridine aryloxyl phosphoramidate ProTide and sofosbuvir, better suppression of viral titers was observed for the former compound. In addition, better protection from ZIKV-induced cytopathic effect was observed for the 2′-*C*-methyluridine aryloxyl phosphoramidate ProTide over sofosbuvir in a highly ZIKV-susceptible stem cell model of infection with strong phenotypic changes during ZIKV infection. These results represent the first broad study of nucleoside analogue ProTides against ZIKV.

## 2. Materials and Methods

### 2.1. Chemistry

All nucleoside analogue ProTides, including McGuigan aryloxy phosphoramidite ProTides and 2-(methylthio)ethyl tryptamine ProTides were synthesized using established phosphorus chemistry [25,26]. The 2′-*C*-ethenyluridine ProTide was synthesized using a known ethynyl ribose precursor 1, which was obtained by following a previously reported synthetic route (Scheme 1) [27]. The ethenyl group of compound **2** was obtained by hydrogenation of the alkyne using 5% Pd/BaSO4 in (1:1) quinoline/benzene without the loss of the benzoyl protecting groups [28]. Ribosylation of uracil to compound **2** using BSA and SnCl4 in ACN afforded benzoyl-protected 2′-*C*-ethenyluridine **3**. The deprotection of compound **3** was performed using NH_3_/MeOH in MeOH to afford 2′-*C*-ethenyluridine 4, which was then treated with *t*BuMgCl in THF followed by *N*-[(*S*)-(2,3,4,5,6-pentafluorophenoxy)phenoxyphosphinyl]-l-alanine 1-methylethyl ester and to afford 2′-*C*-ethenyluridine aryloxy phosphoramidate **5**.

#### 2.1.1. General Synthetic Experimental

All reagents, chemicals, and nucleoside analogues precursors were purchased from Acros Organics, Carbosynth, Chem-Impex International, AK Scientific, and Fisher Chemical at ACS reagent grade or higher purity and used as received. (3R,4R,5R)-5-(benzoyloxy)methyl)-3-ethynyltetrahydrofuran-2,3,4-triyl tribenzoate **1** was synthesized by following a previously reported procedure [27]. Synthetic details for aryloxy phosphoramidate ProTides other than for 2′-*C*-ethynyluridine and 2′-*C*-ethenyluridine were published previously by our group [24]. All reactions were carried out in either an oven-dried round bottom flask or Schlenk tube under a nitrogen atmosphere, using commercially available anhydrous solvents, and monitored by thin-layer chromatography, with detection by UV light. ^1^H NMR and ^31^P NMR spectra were acquired on a Varian 400-MHz spectrometer and recorded at 298 K. Chemical shifts were referenced to the residual protio solvent peak and are given in parts per million (ppm). Splitting patterns are denoted as s (singlet), d (doublet), dd (doublet of doublet), dq (doublet of quartet), ddd (doublet of doublet of doublet), ddt (doublet of doublet of triplet), t (triplet), td (triplet of doublet), tdd (triplet of doublet of doublet), q (quartet), and m (multiplet).

##### 2′-C-ethynyluridine Aryloxy Phosphoramidate

To a stirred suspension of 2′-*C*-ethynyluridine (0.09 mmol, dried under vacuum at 50°C overnight) in dry tetrahydrofuran (THF; 1 ml) was added a 2.0 M solution of tert-butyl magnesium chloride in THF (96 μl, 0.19 mmol). The mixture was stirred at 0 °C for 30 min and then allowed to warm to room temperature and stirred for an additional 30 min. The reaction mixture was then cooled to 0°C and *N*-[(*S*)-(2,3,4,5,6-pentafluorophenoxy)phenoxyphosphinyl]-l-alanine 1-methylethyl ester (46 mg, 0.10 mmol) was added. The reaction mixture was stirred for 18 hours as the temperature was allowed to warm to room temperature. The solvent was removed by rotary evaporation. The reaction mixture was purified first using flash chromatography (a gradient from 0 to 30% methanol in dichloromethane) and then using preparative, normal-phase HPLC (10 to 40% MeOH in dichloromethane gradient) to afford 25.5 mg of the product (54.3%) in **≥** 95% purity. ^1^H NMR (400 MHz, CD_3_OD): δ= 7.65 (d, 1H, *J* = 8.1 Hz), 7.40 - 7.34 (m, 2H), 7.27 (dq, 2H, *J* = 7.7, 1.2 Hz), 7.20 (ddd, 1H, *J* = 8.2, 7.1, 1.1 Hz), 6.04 (s, 1H), 5.61 (d, 1H, *J* = 8.1 Hz), 4.97 (m, 1H), 4.49 (ddd, 1H, *J* = 11.8, 6.0, 2.1 Hz), 4.36 (ddd, 1H, *J* = 11.8, 6.1, 3.7 Hz), 4.16 (d, 1H, *J* = 9.1 Hz), 4.08 (ddt, 1H, *J* = 9.0, 3.9, 2.1 Hz), 3.92 (dq, 1H, *J* = 9.9, 7.1 Hz), 3.08 (s, 1H), 1.35 (dd, 3H, *J* = 7.2, 1.0 Hz), 1.22 (dd, 6H, *J* = 6.3, 1.8 Hz). ^31^P NMR (162 MHz, CD_3_OD) δ= 3.78.

##### 5-((benzoyloxy)methyl)-3-vinyltetrahydrofuran-2,3,4-triyl tribenzoate 2

To a stirring solution of 1 (208.7 mg, 0.362 mmol, 1 equiv.) in benzene (2 mL) and ethanol (2 mL) under H**2** (g) was added 5% palladium on barium sulfate (20.8 mg, 10 wt%) followed by quinolone (22 μL) and was stirred at room temperature for 2 hours. The mixture was diluted in ethyl acetate, washed 3 times with water, and dried over anhydrous sodium sulfate. The reaction mixture was purified using flash chromatography (0 to 30% ethyl acetate in hexane gradient) to afford purified product. (162.7 mg, 0.281 mmol, 77.7%) 1H NMR (400 MHz, CDCl_3_): δ= 8.23 - 8.09 (m, 4H), 8.07 - 8.03 (m, 2H), 7.92 - 7.88 (m, 2H), 7.69 - 7.39 (m, 10H), 7.18 - 7.12 (m, 2H), 6.46 (dd, 1H, *J* = 17.6, 11.2 Hz), 6.25 (d, 1H, *J* = 8.3 Hz), 4.54 (dd, 2H, *J* = 12.2, 4.8 Hz), 4.81 (ddd, 2H, *J* = 8.4, 4.7, 3.9 Hz), 4.73 (dd, 1H, *J* = 12.2, 3.9 Hz), 4.54 (dd, 1H, *J* = 12.2, 4.8 Hz)

##### 5-((benzoyloxy)methyl)-2-(2,4-dioxo-3,4-hihydropyrimidin-1(2H)-yl)-3-vinyltetrahydrofuran-3,4-diyl benzoate 3

Uracil (63.0 mg, 0.562 mmol, 2 equiv.) and 2 (162.7 mg, 0.281 mmol, 1 equiv.) were dried under high vacuum in separate round bottom flasks for 2 hours. Under N_2_ (g) and stirring was added dry acetonitrile (2 mL) to uracil followed by addition of bis(trimethylsilyl)acetamide (550.1 μL, 2.250 mmol, 8 equiv.) Reaction mixture was refluxed at 80°C. for 1 hour then cooled to 0°C. Then compound from step 4 in dry acetonitrile (2 mL) was added to reaction mixture followed by tin (IV) chloride (229.9 μL, 1.968 mmol, 7 equiv.) and heated to 60°C. for 3 hours. Reaction mixture was poured into a separatory funnel containing ice cold water, extracted 3 times with ethyl acetate, and combined organic layer was dried over anhydrous sodium sulfate. The reaction mixture was purified using flash chromatography (0 to 100% ethyl acetate in hexane gradient) to afford purified product. (89.4 mg, 0.153 mmol, 54.6%) 1H NMR (400 MHz, CDCh): δ= 9.22 (s, 1H), 8.09 (m, 4H), 7.86 - 7.82 (m, 2H), 7.63 - 7.56 (m, 2H), 7.63 - 7.56 (m, 6H), 7.29 - 7.21 (m, 2H), 6.65 (s, 1H), 6.12 (dd, 1H, *J* = 17.5, 11.1 Hz), 6.04 (d, 1H, *J* = 5.2 Hz), 5.64 (dd, 1H, *J* = 8.2, 2.1 Hz), 5.46 - 5.40 (dd, 2H), 4.94 (dd, 1H, *J* = 12.3, 3.2 Hz), 4.81 (dd, 1H, *J* = 12.3, 5.7 Hz), 4.66 (td, 1H, *J* = 5.5, 3.2 Hz)

##### 1-(3,4-dikydroxy-5-(kydromethyl)-3-vinyltetrahydrofuran2-yl)pyrimidine-2/4(1H/3H)-dione 4

*Compound **3** (90.5 g, 0.155 mmol, 1 equiv.)* was dried overnight on high vacuum. Under N_2_ (g) was added methanol (1.5 mL) then the reaction mixture was cooled to 0°C followed by dropwise addition of sodium methoxide (86.5 μL, 1.553 mmol, 10 equiv.). Reaction mixture was raised to room temperature and stirred for 1.5 hours. Reaction mixture was cooled to 0°C followed by addition of formic acid until pH=4. Reaction mixture was dried *in vacuo* then purified using flash chromatography (0 to 40% methanol [MeOH] in dicholoromethane gradient) to afford purified product. (30.2 mg, 0.146 mmol, 93.8%) 1H NMR (400 MHz, CDCl_3_): δ= 8.13 (d, 1H, *J* = 8.1 Hz), 5.95 (s, 1H), 5.74 - 5.65 (m, 2H), 5.44 (dd, 1H, *J* = 17.3, 1.3 Hz), 5.26 (dd, 1H, *J* = 10.8, 1.3 Hz), 4.22 (d, 1H, *J* = 9.2 Hz), 4.03 - 3.97 (m, 2H), 3.84 - 3.79 (m, 1H)

##### isopropyl(((5-(2,4-dioxo-3,4-dϊkydropyridimin-1(2H)-yl)-3,4-dϊkydroxy-4-vinyltetrakydrofuran-2-yl)metkoxy)(pkenoxy)pkospkoryl)-L-alaninate 5

To a stirring solution of **4** (36.4 mg, 0.176 mmol, 1 equiv.) in dry tetrahydrofuran (1 mL) at 0°C. was added tert-butyl magnesium chloride (184.5 μL, 0.369 mmol, 2.1 equiv.). Reaction mixture was raised to room temperature and allowed to react for 30 min. Reaction mixture was cooled to 0°C. then *N*-[(*S*)-(2,3,4,5,6-pentafluorophenoxy)phenoxyphosphinyl]-l-alanine 1-methylethyl ester (95.5 mg, 0.211 mmol, 1.2 equiv.) was added and gradually warmed to room temperature overnight. The reaction mixture was purified using flash chromatography (0 to 30% methanol [MeOH] in dichloromethane gradient) to afford purified product. (42.5 mg, 0.0788 mmol, 44.8%) 1H NMR (400 MHz, CD_3_OD): δ= 7.76 (d, 1H, *J* = 8.1 Hz), 7.38 (dd, 2H, *J* = 8.6, 7.2 Hz), 7.31 - 7.25 (m, 2H), 7.25 - 7.16 (m, 2H), 5.94 (s, 1H), 5.68 (dd, 1H, *J* = 17.3, 10.8 Hz), 5.60 (d, 1H, *J* = 8.1 Hz), 5.48 (d, 1H, *J* = 1.4 Hz), 5.44 (d, 1, *J* = 1.4 Hz), 5.27 (dd, 1H, *J* = 10.8, 1.4 Hz), 4.96 (1H, m), 4.58 - 4.49 (m, 1H), 4.45 - 4.38 (m, 1H), 4.17 (s, 2H), 4.00 - 3.87 (m, 1H), 1.38 (dd, 3H, *J* = 7.1, 1.0 Hz), 1.21 (d, 6H, *J* = 6.3 Hz). ^31^P NMR (162 MHz, CD_3_OD) δ= 3.78.

##### (5-(2,4-dioxo-3,4-dikydropyrimidin-1(2H)-yl)-3,4-dikydroxy-4-metkyltetrakydrofuran-2-yl)metkyl (2-(methylthio)ethyl) (2-(1H-indol-3-yl)ethyl)phosphoramidate

2′-*C*-methyluridine (45.3 mg, 0.1790 mmol, 1 equiv.) and triethylammonium 2-(methylthio)ethyl phosphonate (92.1 mg, 0.3579 mmol, 1.5 equiv.) were added to an oven-dried schlenk tube and dried overnight on the high vacuum. To the flask under N**2** (g) was added dry pyridine (2 mL) followed by dropwise addition of trimethylacetyl chloride (60.6 μL, 0.4923 mmol, 2.75 equiv.) and the reaction was stirred for 3 hours at room temperature. The reaction was quenched by saturated sodium bicarbonate and extracted 3 times with dicholormethane. The organic layer was dried over anhydrous sodium sulfate and concentrated in vacuo at room temperature and dried for 2 hours on high vacuum. The dried residue was dissolved in pyridine (2 mL) under N**2** (g). To the reaction mixture was simultaneously added trimethylamine (49.9 μL, 0.3579 mmol, 2 equiv.), tryptamine (114.7 mg, 0.7158 mmol, 4 equiv., dissolved in dry pyridine 2 mL), and carbon tetrachloride (26.0 μL, 0.2685 mmol, 1.5 equiv.) and stirred for 25 min at room temperature. The reaction mixture was diluted with methanol and purified using flash chromatography (0 to 30% methanol in dichloromethane gradient) to afford purified product. (8.0 mg, 0.01443 mmol, 7.3%) ^1^H NMR (400 MHz, CD_3_OD): δ= 7.72 (dd, 1H, *J* = 21.4, 8.2 Hz), 7.55 - 7.50 (m, 1H), 7.33 - 7.29 (m, 1H), 7.06 (t, 2H, *J* = 7.3 Hz), 6.97 (tdd, 1H, *J* = 7.5, 4.6, 2.4 Hz), 5.95 (d, 1H, *J* = Hz), 5.64 (dd, 1H, *J* = 8.1, 3.8 Hz), 4.37 - 4.29 (m, 1H), 4.17 (ddd, 1H, *J* = 11.7, 5.0, 3.1 Hz), 4.04 (dt, 1H, *J* = 10.4, 6.8 Hz), 4.00 - 3.90 (m, 1H), 3.77 (dd, 1H, *J* = 12.4, 9.3 Hz), 3.21 (td, 2H, *J* = 10.9, 5.6 Hz), 2.96 (dt, 2H, *J* = 7.3, 4.4 Hz), 2.66 (td, 2H, *J =* 6.7, 2.3 Hz), 2.07 (d, 3H, *J =* 1.4 Hz), 1.13 (s, 3H). ^31^P NMR (162 MHz, CD_3_OD) δ= 10.52 and 10.41.

**Scheme 1.**
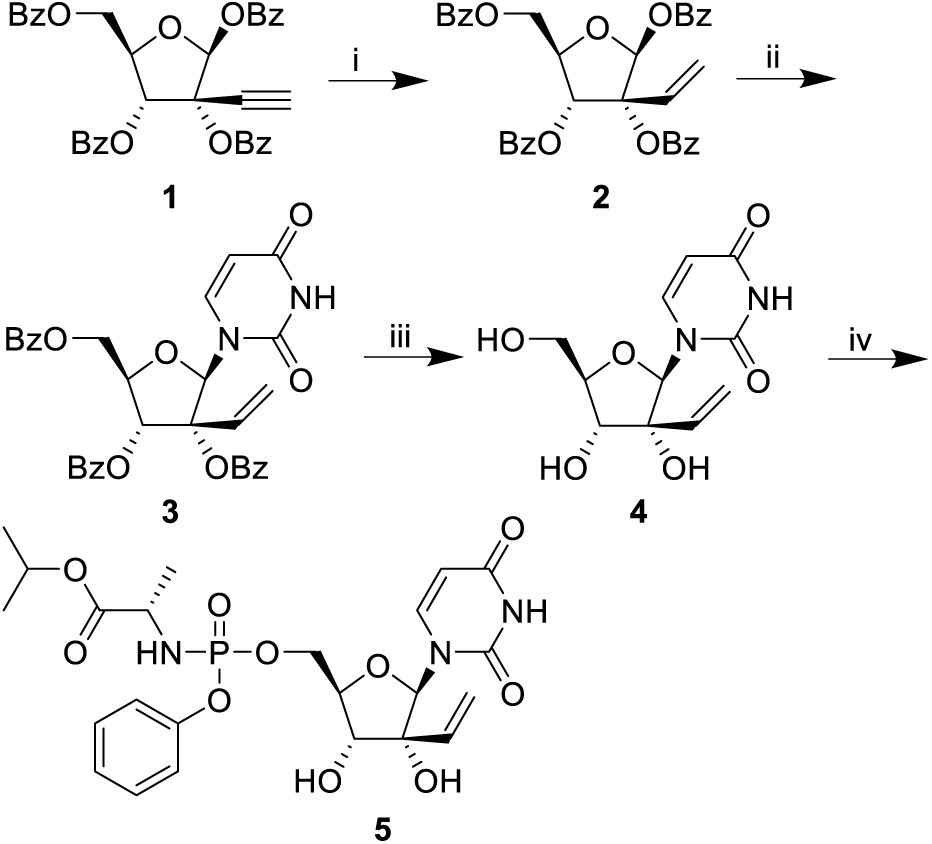
Synthesis of 2′-*C*-ethenyluridine ProTide. i. H2(g), Pd/BaSO4, quinoline, benzene, 78% ii. BSA, uracil, SnCl_4_, ACN, 60 °C to 80 °C, 55**%** iii. NH_3_/MeOH, MeOH, 0 °C to reflux, 94**%**. iv. tBuMgCl, N-[(S)-(2,3,4,5,6-pentafluorophenoxy)phenoxyphosphynyl]-L-alanine 1-methylethyl ester, THF, −70 °C to RT, 45%.

### 2.2. Cell Culture

Human fetal NSCs were purchased from Clontech (human neural cortex; catalog number Y40050). Maintenance of hNSCs was performed in Neurobasal medium without phenol red (Thermo Fisher) and in the presence of B-27 supplement (1:100; catalog number 12587010; Thermo Fisher), N-2 supplement (1:200; catalog number 17502-048; Invitrogen), 20 ng/ml fibroblast growth factor (catalog number 4114-TC-01M; R&D Systems), 20 ng/ml epidermal growth factor (catalog number 236-EG-01M; R&D Systems), GlutaMAX (catalog number 35050061; Thermo Fisher), and sodium pyruvate. Cells were prepared in 6-well plates precoated with laminin (10 mg/ml; catalog number L2020; Sigma) and were grown to near confluence (80 to 90%) prior to passage. For passaging, cells were rinsed gently with phosphate-buffered saline (without calcium and magnesium), and then treated with TryPLE for 5 min at 37°C for cell detachment (catalog number 12563029; Thermo Fisher). The GSC 387 cell line was prepared and maintained similarly to what has been previously described using the media and detachment conditions used for hNSCs described above [29]. Vero cells (ATCC CCL-81) were purchased from ATCC and cultured in Dulbecco′s Modified Eagle Medium (DMEM) (Gibco) with high glucose (4.5g/L), 10% fetal calf serum (FCS) (Sigma), and 1% Penicillin-Streptomycin (PS) (Sigma-Aldrich) at 37°C and 5% CO**2**.

### 2.3. Viruses

Viral strains used in this study were provided through BEI Resources, NIAID, NIH: Zika virus H/PAN/2016/BEI-259634, NR-50210 (GenBank accession number KX198135), Zika virus PRVABC59/Human/2015/Puerto Rico, NR-50240 (GenBank accession number KU501215).

Vero cells were used to expand viral cultures for 2 to 3 serial passages in order to amplify the titers. Centrifugation of infected cell supernatants was performed to remove cell debris and then viral stocks were concentrated through use of a sucrose cushion. Neural maintenance medium base (50% DMEM-F-12 medium-GlutaMAX, 50% Neurobasal-A medium, 1 **×** N-2 supplement, 1 **×** B-27 supplement [Life Technologies]) supplemented with 1**%** DMSO (Sigma) and 5**%** FBS (Gibco) was used to resuspend concentrated viral stocks and aliquots were stored at −80°C. Plaque assay on Vero cells was used to determine viral titers, which were greater than or equal to 2 **×** 10^8^ PFU/ml.

### 2.4. CellTiter-Glo Activity Assay

NSCs (5,000 cells/well) were seeded into 1536-well black, clear bottom plates containing 20-point, 2-fold serial dilutions of ProTides or DMSO vehicle controls, and were either mock-infected, infected with ZIKV H/PAN/2016/BEI-259634 at MOI of 10 or infected with ZIKV PRVABC59/Human/2015/Puerto Rico at MOI of 10. Cell viability was measured using the CellTiter-Glo assay (Promega) 72 hours post-infection. An EnVision plate reader (PerkinElmer) was used for readouts of luminescence intensity. All data were normalized to dimethyl sulfoxide (DMSO) vehicle controls and were expressed as % relative luminescence intensity. Normalized activity and toxicity data were plotted against ProTide concentration and fit to a sigmoidal dose-response curve with variable slope using Prism 6 (GraphPad Software) to generate EC**50** and CC**50** curves.

### 2.5. Bright-field microscopy

GSC 387 cells (5,000 cells/well) were seeded into 1536-well black, clear bottom plates with wells containing either 10 or 30 μΜ sofosbuvir, 10 or 30 μΜ 2′-*C*-methyluridine aryloxyl phosphoramidate ProTide, or DMSO vehicle controls, and were either mock-infected or infected with ZIKV H/PAN/2016/BEI-259634 at MOI of 10. Cell foci images were acquired 72 hours post-infection in bright-field using an ImageXpress Micro automated microscope (Molecular Devices).

### 2.6. Plaque assay

Vero cells were used for the titering experiments for drug-treated and vehicle-treated hNSC cultures infected with ZIKV H/PAN/2016/BEI-259634. Vero cells were seeded at a density of 30,000 cells per well in 96-well plates and incubated in 5% CO_2_ at 37°C in high-glucose DMEM with 1% FBS and 1% penicillin/streptomycin for 24 hours before addition of innocula from ZIKV-infected hNSCs. Inocula were collected from ZIKV H/PAN/2016/BEI-259634-infected NSCs (MOI 0.1) in the presence of 10 μΜ or 30 μΜ 2′-*C*-methyluridine aryloxyl phosphoramidate ProTide, 10 μΜ or 30 μΜ sofosbuvir, mock-infected cultures, infected cultures without compound treatment, and infected cultures with the DMSO vehicle at 48 hours post-infection. Collected inocula were diluted 10-fold in high-glucose DMEM containing 1% FBS and 1% penicillin-streptomycin in 96-well plates. These inocula were added to the pre-prepared Vero cells for 1 hour in a volume of 100 μL and at a temperature of 37°C, then covered with an agarose overlay, and lastly incubated for 72 hours at 37C and 5% CO**2**. Fixation with 4% formaldehyde (final concentration) for 24 hours was then performed for the samples. Following this, the overlay was removed and crystal violet staining was performed to count the ZIKV plaques.

### 2.7. ZIKV RdRP expression and purification

The RdRP domain of ZIKV NS5 (amino acids 2772-3423 of accession number AMB18850) was fused to an *N*-terminal 6x-histidine tag containing a TEV cleavage site, and the cDNA was codon-optimized and synthesized by IDT (San Diego, CA). This construct was then inserted into a pET-17b vector (Invitrogen) and heterologously expressed in the Rosetta strain of *E. coli*. Cultures transfected with the RdRP construct were grown at 37°C in LB media containing 1% ethanol and 100 μg/mL amplicillin until an OD_600_ of 0.6-0.7 was reached. Protein expression was then induced with 100 μM IPTG prior to an additional 5 hours incubation at 20C and lower speed shaking (130 RPM). Cells were then harvested by centrifugation at 9,000 RPM for 30 min at 4°C, and pellets were stored at - 80°C. For purification, pellets were resuspended in lysis buffer (50 mM HEPES at pH 7.5, 20 mM NaCl, 20% glycerol, 0.1% octyl-glucoside, 5 mM β-mercaptoethanol) supplemented with a protease inhibitor tablet (Thermo Fisher). Cells were lysed using a microech) at 4°C. Unbound protein was washed with 50 mE each of the following buffers: wash buffer (50 mM HEPES at pH 7.5, 5 mM β-mercaptoethanol) containing 100 mM NaCl and 10 mM imidazole, wash bfluidizer (Microfluidics International Corporation, Newton, MA) and clarified via centrifugation at 12,000 RPM for 30 min at 4°C. The supernatant was loaded onto TAEON cobalt affinity beads (Clontuffer containing 500 mM NaCl and 20 mM imidazole, and finally wash buffer containing 200 mM NaCl and 30 mM imidazole. Protein was eluted in wash buffer containing 200 mM NaCl and a linear imidazole gradient of 30-300 mM. Protein fractions were pooled and dialyzed overnight against 50 mM HEPES at pH 7.0, 100 mM NaCl, 10% glycerol, 0.01% CHAPS, and 5 mM dithiothreitol at 4°C. Protein was concentrated to **ɥ** 6 μΜ and stored in single-use aliquots at −80°C.

### 2.8. Single nucleotide incorporation assay

To prepare the double-stranded RNA substrate, primer (5’-AGUUGUUGAUC) was ^32^P-labeled at the 5′ position upon incubation with [**γ**-^32^P] ATP (PerkinElmer, Waltham, MA) and T4 polynucleotide kinase (NEB, Ipswich, MA) at 37°C for 30 min. Excess [**γ**-^32^P] ATP was removed with a Bio-Spin 6 column (Bio-rad, CITY). Followed by heat inactivation, the 5′-^32^P labeled primer was then combined with excess template (5’-AUUCACUCAGAUCAACAACU-3′). Primer and template were annealed via step incubation at 95°C (5 min), 55°C (15 min), and 37°C (10 min) to generate primer/template substrate (P/T substrate) where the next correct nucleotide to be incorporated is UTP.

Single nucleotide incororation assays were modeled from previous work [19]. Briefly, ZIKV RdRP (1 μM) and P/T substrate (200 nM) were pre-incubated at 37°C for 20 min. Reactions were initiated at 37°C with the addition of 150 μM UTP or analog in 50 mM HEPES pH 7.5, 15% glycerol, 15 mM NaCl, 7.5 mM MnCl**2**, and 10 mM dithiothreitol. At specified time points, reaction aliquots were quenched with ethylenediamine tetra-acetic acid (EDTA) to a final concentration of 0.3 M.

Prior to loading equi-volume samples on a 20% polyacrylamide, 8 M urea, TBE (89 mM Tris-Borate, 2 mM EDTA) denaturing gel,e quenched samples were combined with dye (formamide, 0.01% xylene cyanol, 0.01% bromophenol blue) and 1 μL of 5 μM D10 (5′-AGTTGTTGAT) trap, followed by incubation at 95°C for 5 min. The trap was used to bind to RNA template to reduce gel smearing [30]. Oligos were separated via electrophoresis at a maximum of 1,900 V and 75 W for 3-6 hours. Phosphorimaging (FLA 9500 imager, GE, Marlborough, MA) was used to visualize the bands, and bands were quantified using ImageQuant software (GE). Plots of percent incorporation versus time were fit to a single exponential equation using Prism (Graphpad Software, San Diego, CA) to compare relative incorporation. While single-turnover conditions are used, non¬physiologically relevant rates were obtained as incorporation occurred during a steady-state time scale. This has been a common issue in the field when measuring ZIKV RdRP incorporation [19.20.31] due to measuring incorporation into a double-stranded template instead of physiologically preferred *de novo* RNA replication (an issue seen in HCV incorporation assays [32]), and perhaps the presence of purification tags [33]. However, since here we were interested in comparing relative incorporation efficiency rather than obtaining rate constants, we and others [19.31] have found this experimental set-up ideal and better suited to scaling up the number of analogues to be assessed.

### 2.9. Structural alignment

A model of ZIKV RdRP in complex with RNA, Mg^2+^ ions, and an incoming ADP modified to an ATP (gc-o3 [34]), generated from ZIKV RdRP stucture PDB 5TFR [35] and the HCV RdRP structure PDB 4WTD [36] was aligned with HCV RdRP [36] using Pymol [37] (160 atoms aligned, RMS = 1.650). Pockets were highlighted by selecting residues 6 Å from the bound nucleotide in the ZIKV (735 atoms selected) and HCV RdRP (598 atoms selected) structures.

## 3. Results

### 3.1. ProTide library design

To investigate the activity of a set of nucleotide analogue ProTides against ZIKV and obtain basic structure-activity relationships, we selected a set of nucleotides designed to inhibit viral RdRPs by three different mechanisms: obligate chain termination, nonobligate chain termination, and lethal mutagenesis (Figure 1) [38,39]. Past work—and the successful results of drug development against HCV—shows that nucleosides alone lack adequate potency, owing to slow phosphorylation in the target cells [40]. McGuigan′s nucleotide aryloxy phosphoramidates include amino acid esters and are well established; sofosbuvir includes this functional group, and we chose it as our starting point [9,41]. One potential drawback of the McGuigan design is that unmasking is initiated by the action of carboxyesterases, which are highly active in the hepatocytes targeted by HCV, but less active in the neural cell targets of ZIKV [40]. For that reason, we chose to investigate a recently developed methylthioethyl tryptamine ProTide from Wagner, which is unmasked by chemical cleavage of methylthioethanol followed by tryptamine removal by HINT1 [26]. HINT1 also catalyzes P-N bond cleavage in the classic McGuigan ProTide. Our hypothesis was that the carboxyesterase-independent unmasking of the methylthioethyl tryptamine ProTide would provide greater potency against ZIKV in human neural stem cells.

**Figure 1.**
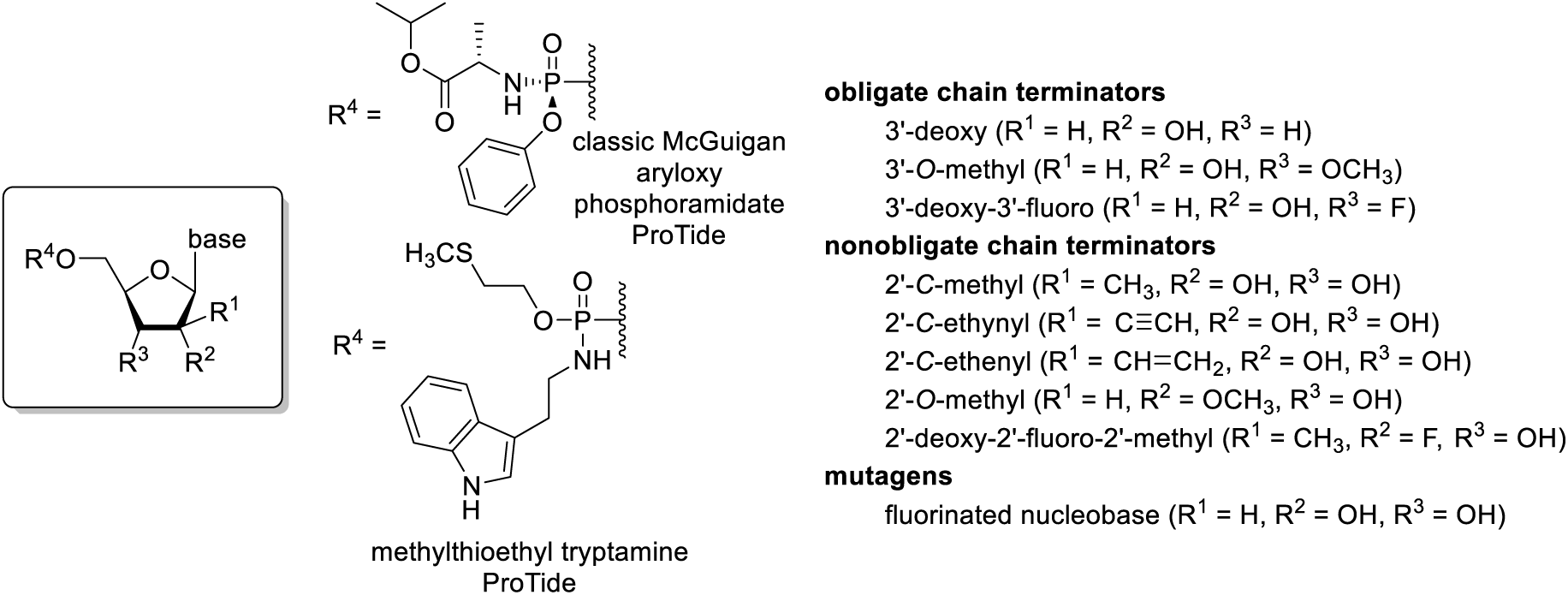
Design and structure of nucleotide analogue inhibitors tested against ZIKV in this study.

The selection of nucleoside analogues for testing focuses on ribose modifications known to induce chain termination *in vitro* when used as a nucleoside triphosphate analogue with the viral RdRP. Some of the selected compounds, e.g. the 3′-deoxy and the 3′-*0*-methyl, are obligate chain terminators, but phosphorylation of this class of compounds in cells is often inefficient [18,19].

Better success in antiviral drug developmentt has been found when using nonobligate terminators, especially 2′ modifications that alter he nucleoside conformation and prevent extension after incorporation during RNA synthesis [14,31]. We include sofosbuvir in this set as a point of reference, along with ProTides of the 2′-*C*-methyl ribosides, which have known potency against the *Flaviviridae* family [19,31,42]. Also included are new ProTides of 2′-*C*-ethynyl- and 2′-*C*-ethenyluridine. 2′-*C*-ethynyl ribosyl triphosphates are known sub-μΜ inhibitors of flavivirus RdRPs [31,43], and we prepared the new 2′-*C*-ethenyluridine ProTide to probe the active site of the RdRP and the metabolic consequences of further modification of the 2*′-C-β* position. A 5-fluorouridine ProTide was included in our set of analogues to test inhibition by lethal mutagenesis, although toxicity of mutagenic nucleoside analogues is typically prohibitive of clinical applications.

### 3.2. Activity against ZIKV and toxicity of nucleoside analogue ProTides in human neural stem cells

To profile the activity and toxicity profiles of our library of ProTides, we screened compounds in a 20-point, 2-fold dose response with highest concentration of 50 μΜ against ZIKV PRVABC59/Human/2015/Puerto Rico in neural stem cells (multiplicity of infection (MOI) of 10) using a luminescent cell-viability assay (CellTiter Glo), and determined EC50 (effective concentration 50) and CC 50 (cytotoxic concentration 50) values from this data (Table 1). We observed that the compounds with the best selectivity indexes (SIs) were the 2′-*C*-methyluridine aryloxyl phosphoramidate ProTide (EC5oPRVABC59: 0.97 μΜ, CC5oPRVABC59: 46.65 μΜ, SI: > 48) and the 2′-*C*-ethynyluridine aryloxyl phosphoramidate ProTide (EC50PRVABC59: 0.97 μM, CC5oPRVABC59: 46.65 μΜ, SI: > 48). These outperformed sofosbuvir in terms of anti-ZIKV activity (EC5oPRVABC59: 35.31 μΜ, CC5oPRVABC59: >50 μΜ, SI: > 1.4). Activity was observed for the 2′-*C*-methyladenosine aryloxyl phosphoramidate ProTide (EC5oPRVABC59: 9.78 μΜ, CC5oPRVABC59: > 50 μΜ, SI: > 5.1) and 2′-*C*-methylcytidine aryloxyl phosphoramidate ProTide (EC5oPRVABC59: 47.8 μΜ, CC5oPRVABC59: > 50 μΜ, SI: > 1.0), albeit at lower levels. None of the other ProTides tested displayed activity. Interestingly, the 2′-*C*-ethenyluridine aryloxyl phosphoramidate ProTide was completely inactive and quite toxic to the host cells, suggesting that this functional group may be more promiscuous in terms of target binding or reactive with cellular components [44]. Replacement of the classic McGuigan ProTide masking group with methylthioethyl tryptamine phosphoramidate for the 2′-*C*-methyluridine ProTide completely abolished activity. This result was not expected because the classic McGuigan ProTides are unmasked with the involvement of both a carboxyesterase and HINT1, whereas the methylthioethyl tryptamine phosphoramidate requires only HINT1 in addition to spontaneous, chemical steps [26]. Clearly, cell line specificity for ProTide strategies must be considered in the design of nucleotide analogue inhibitors of ZIKV.

**Table 1.** Anti-ZIKV activity and toxicity of nucleoside analogue ProTides in neural stem cells.

| **Nucleoside**<br>**Analogue**<br>**ProTide** | **EC<sub>50</sub>**<br>**PRVABC59**<br>**ZIKV, µM** | **CC<sub>50</sub>**<br>**PRVABC59**<br>**ZIKV, µM** | **SI<sup>1</sup>**<br>**PRVABC59**<br>**ZIKV** |
| --- | --- | --- | --- |
| 2'-C-Methylcytidine aryloxyl phosphoramidate | 47.8 +/- 1.1 | > 50 | > 1.0 |
| 3'-Deoxyadenosine aryloxyl phosphoramidate | > 50 | > 50 | > 1.0 |
| 5'-Fluorocytidine aryloxyl phosphoramidate | > 50 | 10.1 +/- 1.1 | > 0.2 |
| 3'-O-Methyluridine aryloxyl phosphoramidate | > 50 | > 50 | > 1.0 |
| 2'-O-Methylcytidine aryloxyl phosphoramidate | > 50 | > 50 | > 1.0 |
| 2'-C-Methyluridine aryloxyl phosphoramidate | 1.0 +/- 1.0 | 46.7 +/- 1.1 | > 48 |
| 2'-C-Methyladenosine aryloxyl phosphoramidate | 9.8 +/- 1.1 | 41.5 +/- 1.2 | > 4.2 |
| 3'-Deoxycytidine<br>aryloxyl<br>phosphoramidate | > 50 | > 50 | > 1.0 |
| 3'-Deoxy-3'-<br>Fluoroguanosine<br>aryloxyl<br>phosphoramidate | > 50 | > 50 | > 1.0 |
| 2'-C-Ethenyluridine<br>aryloxyl<br>phosphoramidate | > 50 | 9.6 +/- 1.1 | < 0.2 |
| 2'-C-Ethynyluridine<br>aryloxyl<br>phosphoramidate | 0.3 +/- 1.1 | > 50 | > 147 |
| 2'-C-Methyluridine 2-<br>(methylthio)ethyl<br>phosphoramidate | > 50 | > 50 | > 1 |
| Sofosbuvir | 35.3 +/- 1.0 | > 50 | > 1.4 |

<sup>1</sup> SI = selectivity index: CC<sub>50</sub>/EC<sub>50</sub>.
<sup>2</sup> Data values are the represented +/- standard error (SE), n=4.

### 3.2. Protection from ZIKV-induced. effect by 2′-C-methyluridine aryloxyl phosphoramidate ProTide and sofosbuvir in a ZIKV-sensitive glioblastoma stem cell line

We and others have previously shown that certain lines of brain tumors with stem cell-like genetic programs, such as glioblastomas, are highly susceptible to ZIKV infection [29,45–47]. These cells display a striking phenotypic change upon infection. To highlight the ability of a representative ProTide hit from our library, 2′-*C*-methyluridine aryloxyl phosphoramidate, to protect against ZIKV-induced cytopathic effect, we treated the glioblastoma stem cell GSC 387 with either 10 or 30 μΜ 2′-*C*-methyluridine aryloxyl phosphoramidate and compared this to 10 or 30 μΜ sofosbuvir or DMSO vehicle control treatment. Samples were prepared in both ZIKV-infected (MOI of 10) and uninfected conditions. After 72 hours post-infection, bright-field microscopy images were obtained for each sample and are shown in Figure 2. While both compounds were able to protect against the cytopathic effect at 30 μΜ (intact cell foci are seen), the 2′-*C*-methyluridine aryloxyl phosphoramidate ProTide was better able to protect against ZIKV-induced cytopathic effect at the lower concentration of 10 μΜ than sofosbuvir, which is in line with our EC_50_ data in neural stem cells. These data suggest that monosubstituted 2′-*C*-modified uridylate aryloxyl phosphoramidate ProTides may be an attractive chemical series to pursue for more potent and selective ZIKV polymerase inhibitors than sofosbuvir.

**Figure 2.**
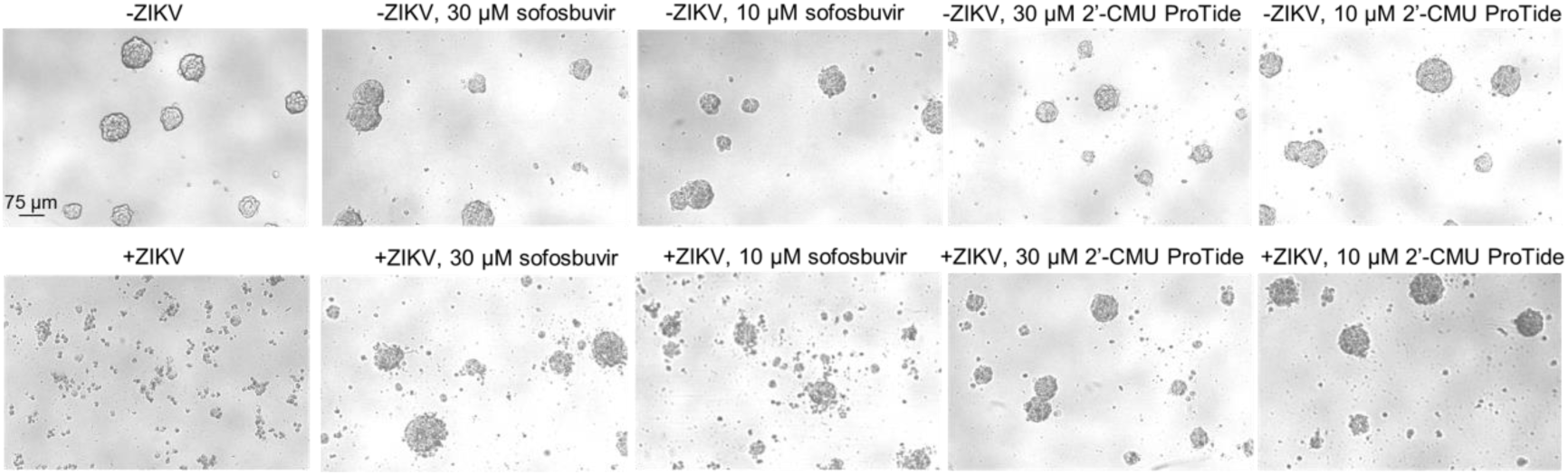
Protection from ZIKV-induced cytopathic effect by 2′-*C*-methyluridine aryloxyl phosphoramidate ProTide and sofosbuvir in a ZIKV-sensitive glioblastoma stem cell model. DMSO-treated GSC 387 cells (-ZIKV and +ZIKV, H/PAN MOI 10), 10 or 30 μΜ sofosbuvir-treated cells or 10 or 30 μM 2′-*C*-methyluridine aryloxyl phosphoramidate ProTide (2′-*C*MU ProTide)-treated cells were incubated for 72 hours at 37C and 5% CO2 and cell foci were subsequently imaged in bright-field.

### 3.3 Repression of ZIKV titers by 2′-C-methyluridine aryloxyl phosphoramidate ProTide and sofosbuvir in neural stem cells

To further assess the antiviral activity of the 2′-*C*-methyluridine aryloxyl phosphoramidate ProTide against ZIKV, we performed plaque assays at 48 hours post-infection to titer ZIKV (H/PAN, MOI of 0.1) levels during treatment with either 10 or 30 μΜ of 2′-*C*-methyluridine aryloxyl phosphoramidate ProTide or 10 or 30 μΜ of sofosbuvir in human fetal neural stem cells (Figure 3). At 30 μΜ compound, both 2′-*C*-methyluridine aryloxyl phosphoramidate ProTide and sofosbuvir were able to repress viral titers to undetectable levels. However, only 2′-*C*-methyluridine aryloxyl phosphoramidate ProTide was able to reduce ZIKV titers to undetectable levels at 10 μΜ compound, again highlighting its superior antiviral activity compared to sofosbuvir.

**Figure 3.**
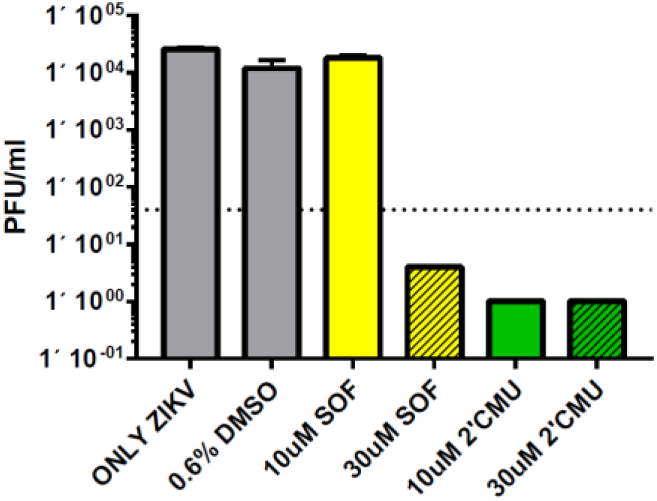
Reduction in ZIKV titers by 2′-C-methyluridine aryloxyl phosphoramidate ProTide and sofosbuvir in neural stem cells as observed by plaque assay. Viral infections were conducted in human fetal neural stem cells (H/PAN ZIKV, MOI 0.1) using vehicle-untreated (ONLY ZIKV), DMSO vehicle-treated, 10 or 30 μΜ sofosbuvir (SOF)-treated, and 10 or 30 μΜ 2′-*C*-methyluridine aryloxyl phosphoramidate ProTide (2′CMU)-treated samples. Virus was titered as described in the methods section and plaque forming units (PFU) per ml were calculated for each sample, +/- standard error (SE), n=4.

### 3.4. Enzymatic incorporation of the active triphosphate metabolites of 2′-C-methyluridine and sofosbuvir over time reveal higher levels of incorporation for the former compound

Single nucleotide incorporation assays were used to compare the relative preference of ZIKV RdRP for UTP versus the UTP analogs 2′-*C*-methyluridine triphosphate and 2′-fluoro-2′-*C*-methyluridine triphosphate (the active form of sofosbuvir) (Figure 4). Both analogues appeared to serve as chain terminators of RdRP polymerization as previously reported [19,31]. Unsurprisingly, UTP was most efficiently incorporated, with 50% incorporation by 5 min and full incoporporation by ~30 min, followed by misincorporating extension. 2′-fluoro-2′-*C*-methyluridine triphosphate had significantly less efficient incorporation by ZIKV RdRP, as previously reported [19,31,48]. Incorporation plateaued at 15%, and half maximal incorporation was only reached after nearly 90 min. In contrast, 2′-*C*-methyluridine triphosphate reached half of its eventual maximal incorporation of 35% after ~45 min, showing improved incorporation over the active form of sofosbuvir. These trends are supported by previous work focusing on biochemical characterization of 2′-*C*-methyluridine triphosphate and 2′-fluoro-2′-*C*-methyluridine triphosphate incorporation [19,20]. Importantly, here we verify the preference of 2′-*C*-methyluridine over sofosbuvir in both biochemical and cell-based assays, suggesting a robust inhibitor testing pipeline that supports the hypothesis that cellular efficacy occurs as a result of RdRP inhibition by nucleoside analogs.

**Figure 4.**
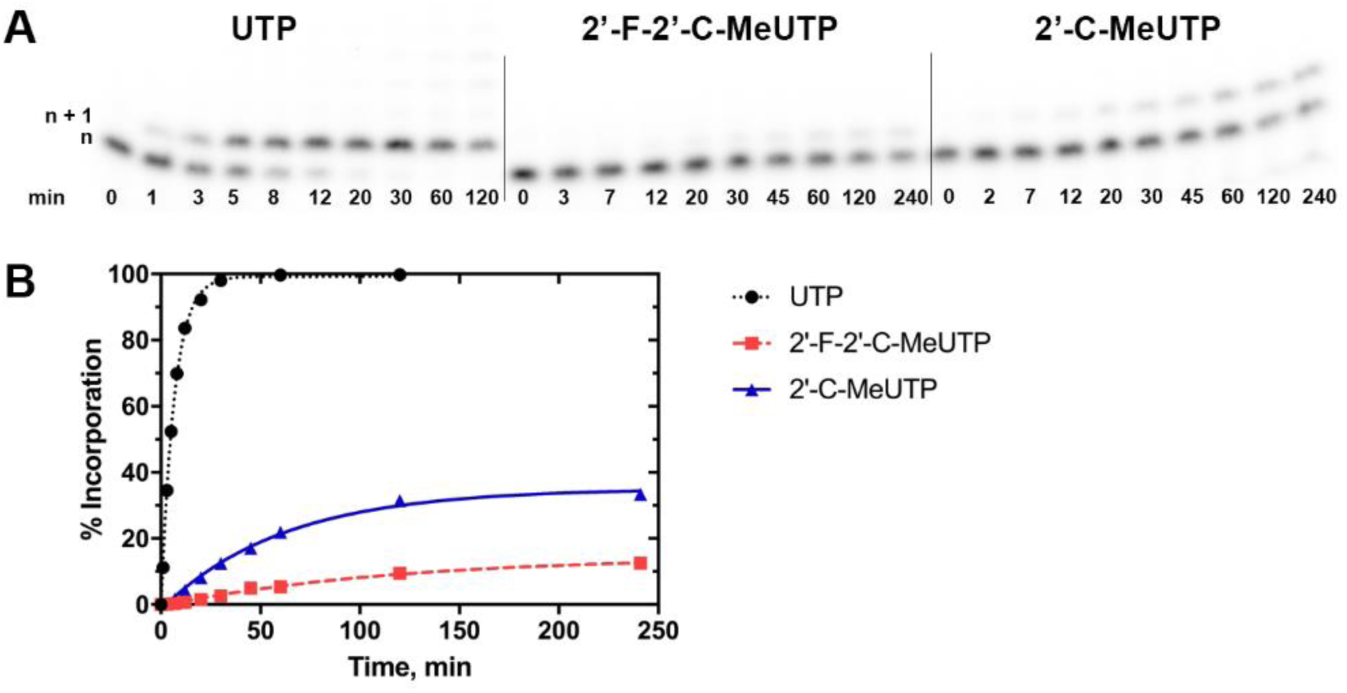
Kinetic characterization of nucleotide and nucleotide analogue incorporation by ZIKV RdRP. (A) ZIKV RdRP and P/T substrate (n) was mixed with excess UTP, 2′-F-2′-*C*-methyluridine triphosphate (2′-F-2′-*C*-MeUTP) or 2-C-methyluridine triphosphate (2′-*C*-MeUTP) and the reaction was quenched at the indicated timepoints. Substrate and the single nucleotide extension product (n + 1) were separated by using a 20% polyacrylamide denaturing gel and bands were quantified to determine percent incorporation and plotted against time. (B) A single exponential fit was used to determine an observed rate of incorporation, *k*_obs_ (**%** incorporation min^−1^) of 0.151 ± 0.004, 0.0085 ± 0.001, and 0.016 ± 0.001, for UTP (dotted black line), 2′-F-2′-*C*-MeUTP (dashed red line), and 2′-*C*-MeUTP (solid blue line), respectively. Standard error is reported as deviation from the fit.

### 3.5. Molecular superpositioning of the ZIKV NS5 active site: consequences for 2′-C-derivatitized nucleoside analogues

A previously prepared model of ZIKV RdRP in complex with RNA and ATP generated from PDB 5TFR [35] and PDB 4WTD [36], gc-o3 [34], was aligned with a crystal structure of HCV RdRP complexed with RNA and sofosbovir diphosphate (PDB 4WTG [36]) (Figure 5). ZIKV RdRP is shown in magenta, and HCV RdRP is shown in green. Of note, HCV D225 swings upward to accommodate the analogue, while in this model from [34], the corresponding D1141 in ZIKV points in 180° in the opposite direction. Interestingly, D1141 in ZIKV was shifted ~90° counterclockwise to overlap with the ribose binding in an apo stucture of ZIKV RdRP (not shown, PDB 5WZ3 [49]). Residues surrounding D1141 in ZIKV are larger than those surrounding the corresponding D225 in HCV, suggesting possible less flexibility to accommodate additional 2′ ribose groups. This may account for the observed activity of the 2′-*C*-ethynyluridine nucleoside for ZIKV and allow for additional modifications at this position for the design of selective ZIKV inhibitors.

**Figure 5.**
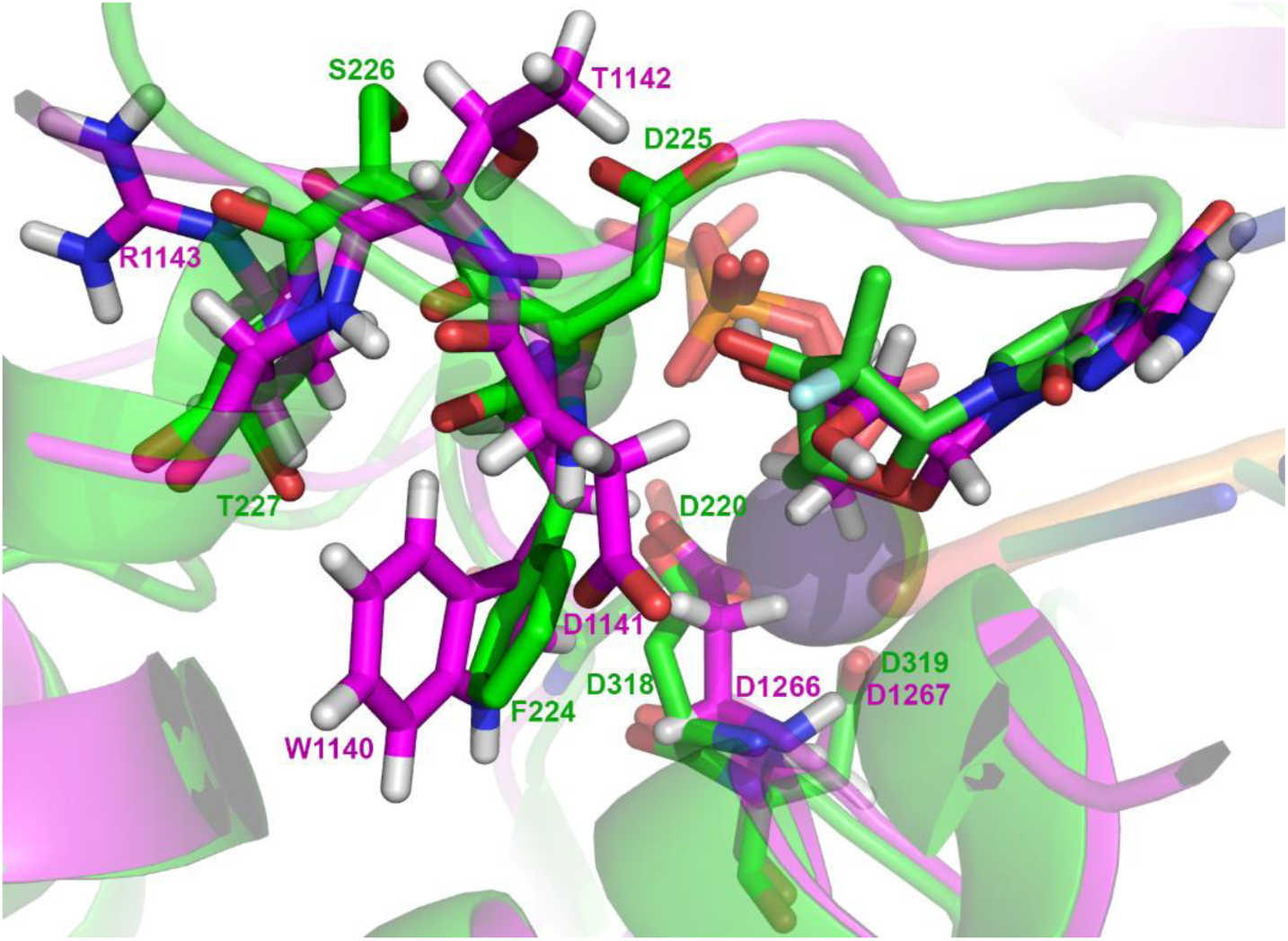
Structural superpositioning of ternary complexes of a ZIKV RdRP model and HCV RdRP crystal structure. Residues highlighted in green are from the ZIKV RdRP model gc-o3 [34] containing the polymerase, RNA, and ATP and those in magenta are from the crystal structure 4WTG containing the HCV RdRP, RNA and sofosbuvir diphosphate [36]. Residues near the 2′-fluoro group in the sugar of sofosbuvir diphosphate are highlighted in stick format in both proteins (R1143, T1142, and W1140 in ZIKV, and T227, S226, and F224). Catalytic residues D220, D318, and D319 in HCV and D1266 and D1267 are also highlighted.

### 4. Discussion

In this work, we have demonstrated that ProTides other than sofosbuvir have promise as potential antiviral drugs to treat ZIKV infections. Interestingly, an apparent bias towards 2′-*C*-modified uridylate ProTides (with the exception of the ethenyl derivative) was observed, suggesting a possible nucleobase bias to compound activity. Recent biochemical data examining levels of off-target incorporation of nucleoside analogue triphosphates by the human mitochondrial RNA polymerase may help explain the phenotypic relationship we observed: both 2′-*C*-methyluridine triphosphate and 2′-*C*-methyl-2′-*C*-fluorouridine triphosphate (the active metabolite of sofosbuvir) show much lower levels of incorporation by this host enzyme than 2′-*C*-modified analogues with other nucleobases.[50]. We propose that this may account for the better safety profile observed for sofosbuvir compared to other 2′-*C*-modified compounds that were tested in clinical trials. This may also suggest that there is greater selectivity for 2′-*C*-modified uridylate incorporation by flavivirus polymerases over host polymerases, though the exact mechanism by which this occurs should be examined in future studies.

We also observed that changing the ProTide functionality of 2′-*C*-methyluridine from an aryloxyl phosphoramidate masking group to a 2-(methylthio)ethyl phosphoramidate resulted in a complete loss of activity, suggesting that the local metabolic environment of the host cell is critical for processing of the ProTide functionality. Further studies examining the relationship between different ProTide functionalities and the cell type dependence of their antiviral activity, as well as testing the impact of these effects using *in vivo* animal model testing are planned in our research groups.

We have demonstrated here for the first time that ProTide technology can be broadly applied to anti-ZIKV drug development with nucleobase and ProTide group selectivity observed. These data will guide future work to design highly selective, safe and bioavailable compounds for the treatment of ZIKV infections.

## Author Contributions

J.A.B., M.C., S.B., G.S.W., L.A.L., A.E.C., Z.Z., C.D.S., B.W.P., and J.L.S.N. conceived and designed the experiments; J.A.B., M.C., S.B., G.S.W., L.A.L., A.E.C., and Z.Z. performed experiments; J.A.B., M.C., S.B., G.S.W., C.D.S., B.W.P., and J.L.S.N. analyzed the data;: J.A.B., M.C., S.B., G.S.W., C.D.S., B.W.P., and J.L.S.N. wrote the original draft of manuscript; J.A.B., M.C., S.B., G.S.W., L.A.L., A.E.C., Z.Z., C.D.S., B.W.P., and J.L.S.N. reviewed and edited the manuscript.

## Funding

This research was supported by Clinical and Translational (CTRI) pilot grant UL1TR001442 (to J.L.S.N.), and a CSUPERB New Investigator grant (to C.D.S). B.W.P., M.C., G.S.W., L.A.L., and C.D.S. thank San Diego State University and the California Metabolic Research Foundation for financial support.

## Conflicts of Interest

The authors declare no conflict of interest. The funders had no role in the design of the study; in the collection, analyses, or interpretation of data; in the writing of the manuscript, or in the decision to publish the results.

